# Genome-Wide Selection Signatures in Nili-Ravi Buffalo (Bubalus bubalis) Reveal a T-Cell Costimulatory and Cytokine-Signaling Gene Network Distinct from Classical Bovine Tuberculosis Candidate Genes

**DOI:** 10.64898/2026.08.10.743898

**Authors:** Atiq Ahmad, Abu bakar, Sheikh Muhammad Laeeque, Waqas Ahmad Khan, Haiba Kaul, Abdul Manan, Hamid Mustafa

## Abstract

Genomic signatures of selection can reveal loci underlying adaptation and disease resistance in livestock populations, but such analyses in water buffalo (Bubalus bubalis) have historically been constrained by the absence of a chromosome-level, species-native reference genome for SNP array data. We re-analyzed genotype data from 85 Nili-Ravi buffalo (Axiom Buffalo Genotyping 90K array, originally positioned using bovine (Bos taurus, UMD3.1) proxy coordinates, by performing a full coordinate liftover to the buffalo-native UOA_WB_1 assembly using an independently published SNP remapping resource. Following quality control (51,209 markers retained), haplotype phasing, and genome-wide integrated haplotype score (iHS) and Wright’s Fst (case/control) selection scans, we evaluated 14 classical bovine-tuberculosis (bTB) candidate genes and identified six additional genes with putative immune function through an unbiased genome-wide screen. None of the 14 classical candidates (including SLC11A1, the Toll-like receptors, and IFNG) reached genome-wide significance in either scan. In contrast, six novel loci TNFSF18, IL2RB, TNFRSF19, IRF2, IL15, and CD28 showed significant iHS or Fst signals, four of which (TNFSF18, IL2RB, IL15, CD28) converge functionally on T-cell costimulation and cytokine receptor signaling (KEGG pathways map04660 and map04060, Bos taurus proxy annotation). Using extended haplotype homozygosity (EHH) decay, haplotype furcation structure, and per-marker haplotype counts as three independent lines of corroborating evidence, we classified these six genes into confidence tiers: TNFSF18 and IL2RB showed the strongest, most balanced support, while CD28 and IL15 signals were driven by very few haplotypes (3 and 5 of 30, respectively) and should be interpreted cautiously pending replication. These findings suggest that adaptive, cell-mediated immune signaling rather than the innate/macrophage-centred mechanisms emphasized by existing bTB candidate gene panels may be a more productive avenue for future selection studies in Nili-Ravi buffalo, while underscoring the value of buffalo-native coordinate systems for accurate genomic inference in this species.

## 1. Introduction

The water buffalo (Bubalus bubalis) is a livestock species of major economic importance across South and Southeast Asia, the Middle East, and the Mediterranean, providing milk, meat, and draught power for millions of smallholder producers. The Nili-Ravi breed, indigenous to the Punjab region of Pakistan and India, is among the highest-yielding riverine buffalo breeds and a focus of ongoing genetic improvement efforts. Despite its agricultural importance, genomic resources for buffalo have historically lagged behind those of cattle, sheep, and goats, constraining the resolution of selection-scan and genome-wide association studies in this species.

A key limitation has been the reference coordinate system used for SNP genotyping arrays. The Axiom Buffalo Genotyping 90K array (Iamartino et al., 2017) was designed prior to the availability of a chromosome-level buffalo genome assembly, and consequently, SNP marker positions were assigned using the bovine (Bos taurus, UMD3.1) reference genome as a proxy. This practice has been widely adopted in subsequent buffalo genomic studies (Mokhber et al., 2019; El-Halawany et al., 2017), and while defensible given the resources available at the time, it introduces a systematic coordinate mismatch: genes and regulatory elements annotated on the bovine genome do not necessarily correspond to their true positions in the buffalo genome, given approximately 5 million years of divergence between the two lineages and documented chromosomal rearrangements between cattle and buffalo karyotypes.

The release of a chromosome-level water buffalo genome assembly (UOA_WB_1; Low et al., 2019) and a subsequent probe-level remapping of the Axiom Buffalo 90K array onto this assembly (Nascimento et al., 2021) now make it possible to reanalyze existing buffalo SNP array datasets using buffalo-native genomic coordinates, correcting for the bovine-proxy artifact inherent in earlier work.

Bovine tuberculosis (bTB), caused by Mycobacterium bovis, remains an important disease concern in buffalo production systems, with variation in host resistance well documented in cattle and increasingly studied in buffalo. A panel of classical candidate genes drawn primarily from cattle bTB genetics studies and including SLC11A1 (NRAMP1), BOLA-DRB3, the Toll-like receptors TLR1/TLR2/TLR4/TLR6, IFNG, NOS2, SP110, VDR, GBP2/GBP6, and CXCR1/CXCR2 represents the prevailing hypothesis-driven framework for investigating host genetic resistance to bTB in Bovinae. Whether this cattle-derived candidate panel captures the relevant genetic architecture of bTB resistance in buffalo, a distinct genus with its own selective history, has not been directly tested using buffalo-native genomic coordinates.

In this study, we re-analyzed Axiom Buffalo 90K genotype data from 85 Nili-Ravi buffalo with a binary phenotype classification, first performing a complete coordinate liftover from the bovine-proxy system to the buffalo-native UOA_WB_1 assembly, then conducting genome-wide integrated haplotype score (iHS) and case/control Fst selection scans. We evaluated the 14 classical bTB candidate genes for evidence of selection and, in parallel, conducted an unbiased genome-wide screen for additional genes with putative immune function. We further assessed the robustness of novel candidate loci using three independent haplotype-based lines of evidence EHH decay, haplotype furcation structure, and per-marker haplotype counts and situated candidate genes within KEGG pathway context using the Bos taurus ortholog database, given the current absence of a functionally annotated buffalo-specific pathway resource.

## 2. Materials and Methods

### 2.1 Animals and genotype data

Genotype data were used in this study was retrieve from the previous described (Atiq et al., 2026). A binary phenotype classification (cases, controls) as described previously.

### 2.2 Coordinate system verification and buffalo-native liftover

Initial inspection of the SNP marker coordinates in the working genotype file revealed that chromosome/position assignments matched the bovine UMD3.1 assembly (Bos *taurus*) almost exactly on a per-chromosome basis (e.g., chromosome 1 maximum marker position of 158,317,029 bp versus a UMD3.1 chromosome 1 length of 158,337,067 bp, a 99.99% match), rather than any buffalo-native assembly. This confirmed that the original marker positions reflected the bovine-proxy coordinate system used in the original Axiom array design, consistent with standard practice for this array (Mokhber et al., 2019; El-Halawany et al., 2017).

Marker coordinates were then lifted over to the chromosome-level buffalo-native UOA_WB_1 assembly (Low et al., 2019) using the updated Axiom Buffalo 90K SNP map published by Nascimento et al. (2021), which re-aligned all array probe sequences directly against UOA_WB_1. Of 56,845 markers, 55,205 (97.1%) received valid buffalo-native coordinates spanning 25 chromosomes (24 autosomes plus the X chromosome), consistent with the true river buffalo karyotype; the remaining 1,640 unmapped markers were excluded from further analysis.

### 2.3 Quality control

Although the samples panel had undergone quality control at the level of the full multi-population dataset, this does not guarantee marker polymorphism within the selected sample Nili-Ravi subset specifically. An empirical check found 3,790 of 55,205 markers (6.9%) were monomorphic within the Nili-Ravi subset despite passing panel-wide filtering; these markers, together with an additional 206 markers exceeding 10% missingness, were excluded. The resulting quality-controlled dataset comprised 51,209 markers across all samples (genotyping rate 99.87%).

### 2.4 Haplotype phasing

Genotypes were converted to VCF format, sorted by chromosome and position, and phased using Beagle (Browning & Browning 2007) with default parameters, producing 51,209 phased markers across all 25 buffalo-native chromosomes.

### 2.5 Genome-wide selection scans

Extended haplotype homozygosity-based integrated haplotype score (iHS) statistics were calculated to identify candidate genomic regions showing evidence of recent positive selection, following the approach described by Voight et al. (2006). The analysis was implemented in the **rehh** R package version 3.2.3 (Gautier and Vitalis, 2012; Gautier et al., 2017). Phased haplotypes were imported separately for each chromosome using data2haplohh() with unpolarized alleles and vcf_reader = “data.table”. Because ancestral alleles were not available, allele polarization was not assumed and selection signals were evaluated using the absolute iHS values. Integrated haplotype homozygosity was calculated with scan_hh(), and standardized iHS values were obtained using ihh2ihs().

Markers with a minor allele frequency (MAF) ≤0.05 were excluded using the min_maf = 0.05 filtering criterion implemented in **rehh** (Gautier et al., 2017). Of the 51,209 markers initially analyzed, 720 were removed, leaving 50,489 markers for the genome-wide iHS scan. The iHS statistic was standardized according to the allele-frequency distribution of the markers, as recommended for iHS-based analyses (Voight et al., 2006; Gautier and Vitalis, 2012). Markers with —log_10_ (p_iHS_) _ 4, corresponding to p_iHS_≤ 1 X 10^—4^, were considered candidate selection outliers. Here, the two-sided iHS p-value was calculated from the approximately standard normal distribution of the standardized iHS statistic as p_iHS_ = 2Φ(—I iHS I)(Gautier et al., 2017).

Population differentiation between cases and controls was assessed using the Weir and Cockerham (1984) estimator of *F*_ST_. The analysis was performed in PLINK v1.9 (Chang et al., 2015) using the --fst case-control option, with the phenotype status defining the two comparison groups. Of the 51,209 markers examined, 49,261 yielded valid *F*_ST_estimates. The mean, or unweighted global, *F*_ST_was 0.00945, whereas the weighted global *F*_ST_was 0.0156.

### 2.6 Gene annotation

Candidate genes were identified using the RefSeq annotation associated with the water buffalo reference assembly **GCF_003121395.1 (UOA_WB_1)**. This assembly corresponds to a river buffalo genome and contains chromosome-and scaffold-level genomic features that can be linked directly to the corresponding RefSeq sequence accessions (NCBI, 2018). The chromosome-to-RefSeq accession correspondence was therefore established from the region features provided in the annotation file itself, rather than from an external chromosome-name conversion table. The annotation comprised 29,572 annotated genes.

Significant SNPs were annotated in R using the **GenomicRanges** package from Bioconductor, which provides tools for representing genomic intervals and performing overlap and nearest-neighbour analyses (Lawrence et al., 2013). Each significant SNP was represented as a single-base genomic interval and compared with the RefSeq gene coordinates. A SNP was classified as **intragenic** when its genomic position overlapped the annotated boundaries of at least one gene. SNPs that did not overlap a gene were assigned to the nearest annotated gene using the nearest() function in **GenomicRanges**. For SNPs overlapping multiple gene features, all overlapping genes were retained where applicable; otherwise, the closest gene was reported as the putative candidate gene.

### 2.7 Classical bTB candidate gene evaluation

A targeted panel of 14 genes was evaluated at the corresponding water-buffalo genomic coordinates to determine whether previously reported bovine tuberculosis (bTB) candidate regions also showed evidence of selection in buffalo. The panel included SLC11A1, BoLA-DRB3, TLR1, TLR2, TLR4, TLR6, IFNG, NOS2, SP110, VDR, GBP2, GBP6, CXCR1, and CXCR2. These genes were selected because they participate in macrophage antimicrobial activity, antigen presentation, innate immune recognition, interferon-mediated responses, nitric-oxide production, intracellular pathogen control, and leukocyte recruitment. Previous studies have reported associations between variation in several of these genes and bTB susceptibility or resistance in cattle, although the strength and type of evidence differ among genes (Allen et al., 2010; Kadarmideen et al., 2011; Sun et al., 2012; Cheng et al., 2016; Chai et al., 2021) (See ***Appendix***).

For each gene, the buffalo-native RefSeq coordinates were obtained from assembly GCF_003121395.1 (UOA_WB_1). Significant SNPs identified in the genome-wide selection scans were intersected with the coordinates of the 14 candidate genes. A SNP was considered intragenic when it overlapped an annotated gene boundary; significant SNPs located outside gene boundaries were assigned to the nearest candidate gene when they fell within the predefined annotation interval. The resulting candidate-gene list was then compared with published cattle and buffalo evidence, while avoiding the assumption that a signal in buffalo necessarily represents the same functional association previously reported in cattle.

### 2.8 Novel immune-gene screening

In addition to the predefined classical bTB candidate-gene panel, all genome-wide significant iHS and F_ST_markers were screened against a broader, independently defined set of immune-related genes. The screening set included genes encoding Toll-like receptors, interleukins and interleukin receptors, interferons and interferon-regulatory factors, chemokines and chemokine receptors, tumour-necrosis-factor superfamily members, complement proteins, major histocompatibility-complex genes, *BoLA* loci, immunoglobulin genes, and associated intracellular signalling adaptors. These gene groups were selected because they represent major components of innate and adaptive immune recognition, inflammatory signalling, antigen presentation, antibody-mediated immunity, and host defence against infectious diseases (Akira et al., 2001; Takeda and Akira, 2004; Medzhitov, 2001).

The screening was performed using regular-expression patterns matched against official gene symbols and available gene names in the buffalo RefSeq annotation. Thus, the procedure was a gene-symbol-based prioritization screen, rather than a formal Gene Ontology (GO) enrichment analysis. This distinction is important because a pattern match can identify genes with immune-related nomenclature but cannot, by itself, establish their functional classification. The biological relevance of this approach is supported by previous buffalo studies describing variation or expression of immune genes, including Toll-like receptors and immunoglobulin loci (Alfano et al., 2014; Verma et al., 2008; Yang et al., 2025).

Because the current *Bubalus bubalis* RefSeq annotation used in this study did not provide GO terms for the annotated genes, GO-based enrichment analysis was not attempted. Consequently, genes identified through the pattern-based screen were treated as putative immune-related candidates requiring further literature-based and functional validation, rather than as confirmed immune genes. In particular, a flagged gene was considered a candidate only when its annotation, gene symbol, genomic position, and available species-specific literature were consistent with a plausible immune function.

### 2.9 Pathway annotation

Pathway annotation was performed by mapping the candidate genes to their corresponding cattle orthologs in the Kyoto Encyclopedia of Genes and Genomes (KEGG). The *Bos taurus* KEGG organism entry, identified by the code **bta**, was used because it provided substantially greater gene-symbol and pathway coverage for the candidate loci than the buffalo entry available at the time of analysis. Although KEGG includes a *Bubalus bubalis* organism entry (**bbub**), the retrieved buffalo record, identified as T05912, contained only 15 gene models, none of which could be reliably matched to recognizable gene symbols in the present dataset. Consequently, the native buffalo entry was not considered sufficiently informative for pathway-level interpretation in this analysis. KEGG pathway membership was therefore inferred from one-to-one or high-confidence *Bubalus bubalis–Bos taurus* ortholog relationships, rather than from direct species-specific buffalo annotations (Kanehisa et al., 2021; Kanehisa et al., 2023).

As an independent assessment of annotation completeness, the candidate genes were also analyzed using the Database for Annotation, Visualization and Integrated Discovery (DAVID). When *Bubalus bubalis* was selected as the background species, annotation coverage was very limited, with only 1 of 5 genes receiving an annotation in the evaluated functional categories. In contrast, use of *Bos indicus* as a closely related bovine proxy increased the coverage to as many as 6 of 6 genes. This comparison supported the use of bovine orthologs for exploratory pathway annotation, but the coverage difference was interpreted as evidence of database annotation bias rather than biological evidence that cattle and buffalo possess different pathway functions (Huang et al., 2009; Dennis et al., 2003).

The resulting KEGG pathways should therefore be interpreted as orthology-based functional predictions. Pathway assignments do not demonstrate that the corresponding genes are functionally active in buffalo, nor do they establish a causal relationship with bTB resistance or susceptibility. Final interpretation was restricted to pathways supported by conserved orthology, consistent gene nomenclature, and relevant cattle or buffalo literature. Because both KEGG and DAVID annotations are periodically updated, the database release or access date should be recorded to ensure reproducibility.

### 2.10 Haplotype-based robustness assessment

To evaluate the robustness of the six newly identified immune-associated signals, extended haplotype homozygosity (EHH) decay and haplotype-furcation patterns were examined around the peak SNP for each gene. EHH was calculated using calc_ehh(), and furcation structures were generated with calc_furcation() in the **rehh** package. These procedures characterize the persistence and breakdown of haplotype sharing around a focal marker and provide complementary visual evidence for recent selection signals (Sabeti et al., 2002; Gautier & Vitalis, 2012; Gautier et al., 2017).

For every focal SNP, the numbers of phased haplotypes carrying the major and minor alleles were recorded. Because the available sample was relatively small, the interpretation focused on concordance among three features: the shape of the EHH decay curves, the organization of the furcation trees, and the distribution of haplotypes between the two alleles at the focal marker.

A study-defined three-tier classification was used to summarize the strength of the supporting haplotype evidence. Tier 1 (strong support) required a clear and extended EHH pattern, well-resolved furcation structures for both allele-specific haplotype groups, and a minor allele represented by at least 10 of the 30 haplotypes. Tier 2 (moderate support) was assigned when the EHH, furcation, and haplotype-frequency patterns were broadly consistent but less distinct or incomplete. Tier 3 (suggestive support) included signals with visually pronounced haplotype structure but limited representation of the minor allele five or fewer of the 30 haplotypes. Such patterns were interpreted cautiously because they may reflect haplotype sharing or recent identity-by-descent among a small number of individuals rather than a population-wide selective sweep (Sabeti et al., 2002; Ralph & Coop, 2010).

All analyses were conducted in R version 4.6.0 using rehh version 3.2.3, GenomicRanges, rtracklayer, and qqman, with complementary genotype and marker-level processing performed in PLINK v1.9. The Tier 1–3 categories were used as qualitative robustness descriptors and were not treated as formal statistical significance levels. In addition, furcation diagrams and EHH profiles were interpreted alongside the genome-wide iHS and *F*_ST_results rather than as independent proof of selection.

## 3. Results

### 3.1 Buffalo-native liftover and quality control

The 85 Nili-Ravi buffalo samples contained genotypes for 56,845 array markers. Probe sequences were remapped to the chromosome-level *Bubalus bubalis* reference assembly UOA_WB_1 using the buffalo array resource developed by Nascimento et al. (2021). This procedure successfully assigned buffalo-native coordinates to 55,205 markers, corresponding to a mapping rate of 97.1%.

The remapping converted marker positions from the original cattle-based coordinate system to the chromosome structure of river buffalo. The UOA_WB_1 assembly represents the river buffalo genome with 25 chromosome-level sequences, consistent with the river buffalo diploid karyotype of 2n = 50(Liu et al., 2019). Thus, the analysis used buffalo-specific chromosome coordinates rather than the 30-autosome cattle coordinate framework associated with the original array design.

Quality control was subsequently performed within the Nili-Ravi dataset. Markers that were monomorphic or had an insufficient genotype call rate were removed. After filtering, 51,209 markers remained for downstream population-genetic, selection-scan, and candidate-gene analyses. Therefore, 3,996 mapped markers were excluded during quality control, while the final dataset retained approximately 92.8% of the successfully remapped markers (Figure 1).

**Figure 1:**
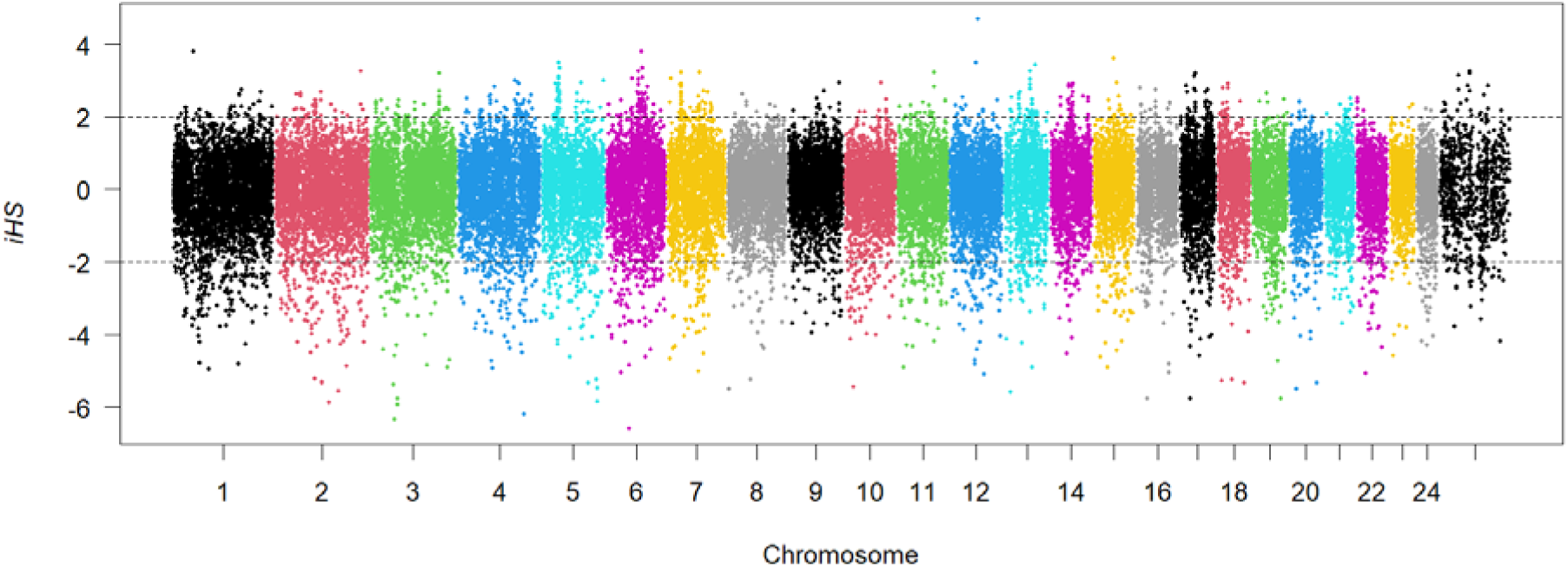
Genome-wide integrated haplotype score (iHS) scan of 15 Nili-Ravi buffalo on buffalo-native (UOA_WB_1) coordinates. Points are colored by chromosome (1–25, including the X chromosome); dashed horizontal lines indicate the standard rehh significance thresholds (|iHS| corresponding to ±2).

### 3.2 Genome-wide iHS and Fst scans

The genome-wide iHS analysis was conducted using buffalo-native marker coordinates and phased haplotypes. A total of 135 markers exceeded the predefined candidate-outlier threshold of —log_10_(p_iHS_) _ 4and were distributed across the buffalo genome (Figure 1). The most prominent signal was detected on chromosome 6 at position 45,435,944 bp, where the iHS-derived value reached —log_10_(p_iHS_) = 10.35. The iHS statistic detects unusually long haplotypes by comparing the decay of extended haplotype homozygosity around the two alleles at a focal marker (Sabeti et al., 2002; Voight et al., 2006; Gautier & Vitalis, 2012). Because the threshold was used to identify candidate selection outliers, these markers were retained for subsequent gene annotation and functional prioritization rather than interpreted as independently confirmed selective sweeps.

The case–control population-differentiation analysis produced a mean *F*_ST_of 0.00945 and a weighted global *F*_ST_of 0.0156. These values were highly similar to those obtained from the original cattle-proxy coordinate dataset, which showed a mean *F*_ST_of 0.0099 and a weighted global estimate of 0.0157. PLINK calculates locus-specific *F*_ST_values using the Weir–Cockerham estimator and reports both raw and weighted global means (Weir & Cockerham, 1984; Chang et al., 2015). The close agreement between the two coordinate systems indicates that remapping the markers to buffalo-native coordinates did not materially change the case–control differentiation results and supports the technical consistency of the coordinate-conversion procedure.

### 3.3 Classical bTB candidate gene panel

None of the 14 predefined classical bTB candidate genes showed evidence of exceeding the selection thresholds in the buffalo-native coordinate analysis. The evaluated genes were *SLC11A1, BoLA-DRB3, TLR1, TLR2, TLR4, TLR6, IFNG, NOS2, SP110, VDR, GBP2, GBP6, CXCR1,* and *CXCR2*. No marker located within or assigned to these genes reached the iHS candidate-outlier threshold of —log_10_(p_iHS_) _ 4, and none fell within the upper 5% of the genome-wide *F*_ST_distribution.

These findings indicate that the selected loci did not show strong evidence of recent haplotype-based selection or pronounced case–control allele-frequency differentiation in the present Nili-Ravi dataset. This result does not exclude their involvement in bTB susceptibility or resistance. Previous studies have reported associations between some of these genes and mycobacterial infection, immune-response phenotypes, or bTB-related traits in cattle and buffalo, but such associations may be population-specific and may not produce detectable selection or differentiation signals in every dataset (Allen et al., 2010; Alfano et al., 2014; Cheng et al., 2016). The absence of a signal in the present analysis may also reflect the modest sample size, the density and distribution of array markers, the allele-frequency spectrum, or differences between historical selection and contemporary disease-associated variation.

Accordingly, these 14 genes were retained as biologically relevant reference candidates but were not classified as selection-supported loci in this study. Their lack of significance also provides a useful contrast with the novel immune-associated genes identified through the genome-wide screening and pathway-prioritization analyses.

### 3.4 Novel immune-associated genes

The genome-wide immune-gene screen identified six genes that were not included in the predefined classical bTB candidate panel: *TNFSF18* on chromosome 5, *IL2RB* on chromosome 4, *TNFRSF19* on chromosome 13, *IRF2* on chromosome 1, *IL15* on chromosome 17, and *CD28* on chromosome 2 (Table 1; Figures 2–3). Five genes were identified through the *F*_ST_analysis, whereas *CD28* was detected through the iHS analysis. None of the six genes was supported by both statistics, and none overlapped the 14 classical bTB candidate genes. These loci were therefore considered novel immune-associated candidates identified through complementary genome-wide selection and population-differentiation analyses.

**Figure 2:**
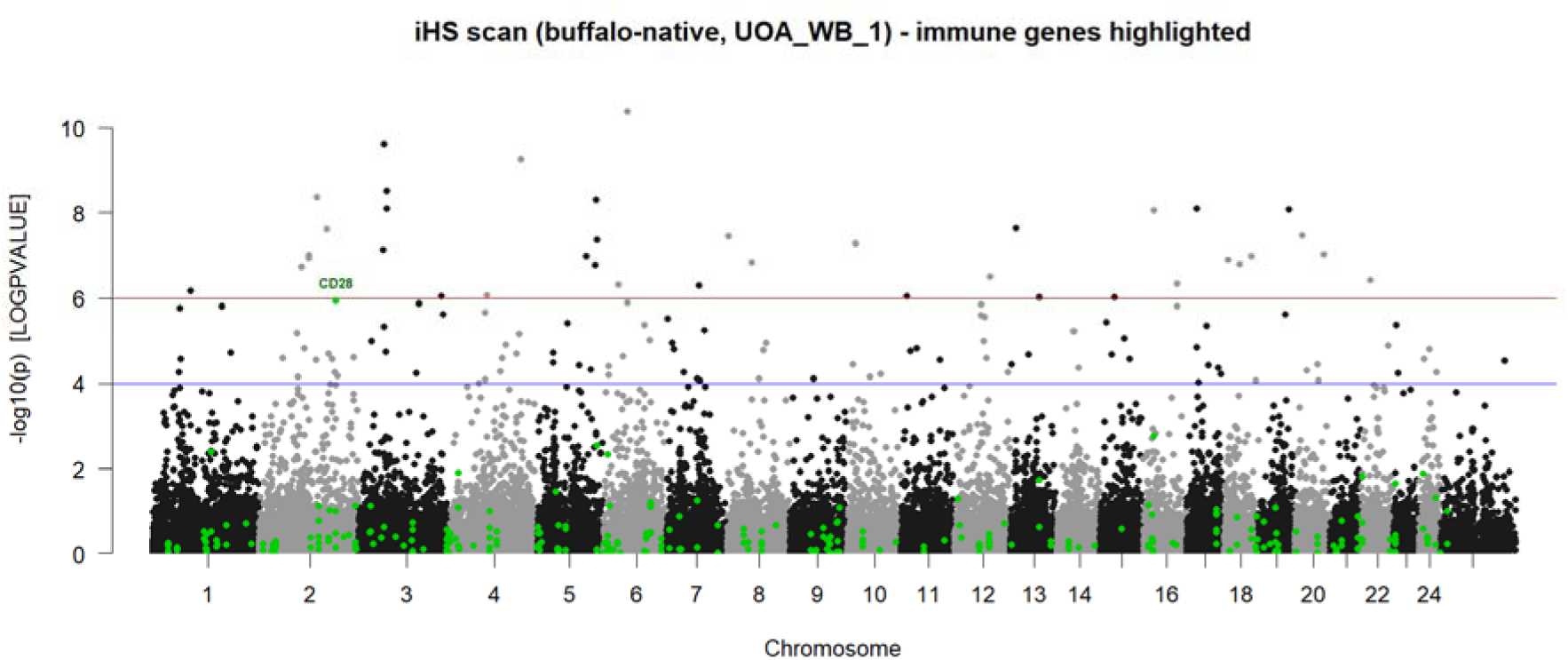
Genome-wide iHS scan with immune-associated genes highlighted (green) and labeled were significant. Blue line: suggestive threshold (−log□□p = 4); red line: genome-wide threshold (−log□□p = 6). CD28 (chromosome 2) is the sole immune-flagged gene reaching the plotted thresholds in this scan.

**Figure 3:**
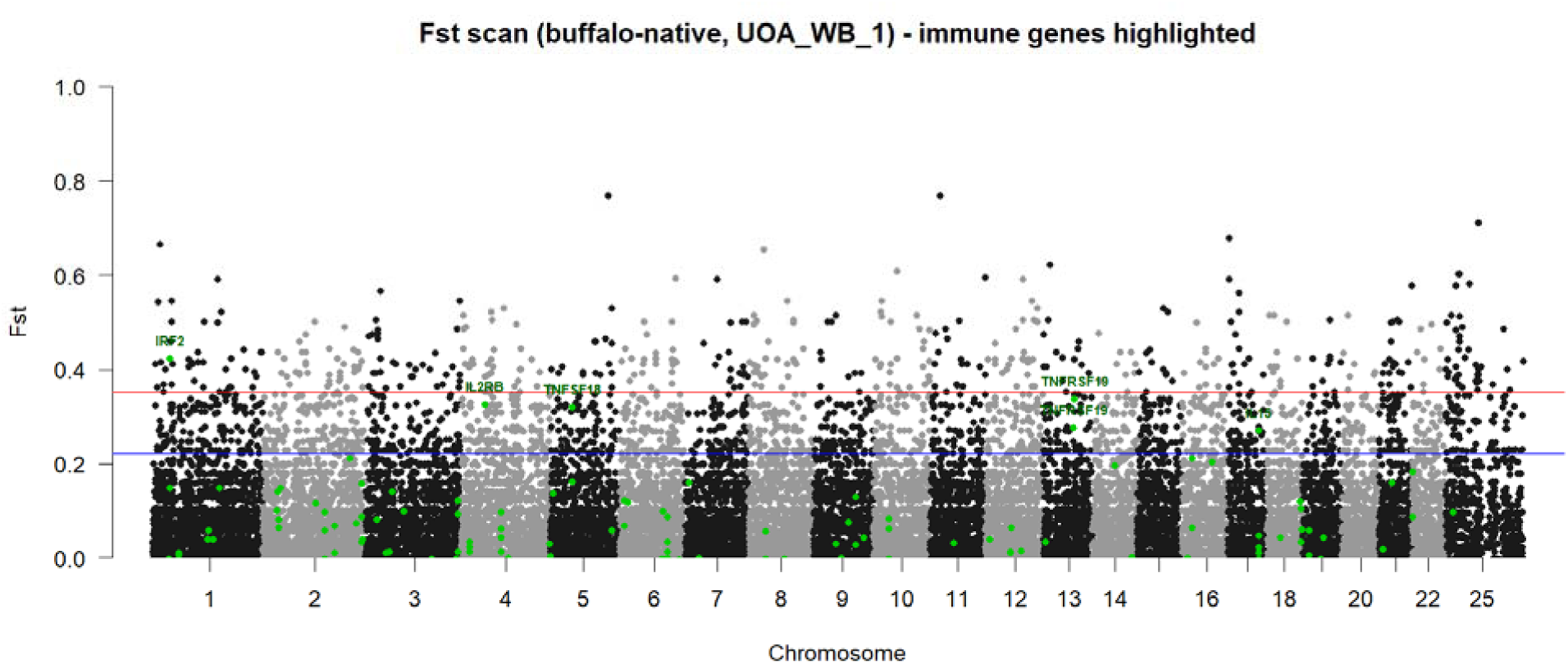
Genome-wide case/control Fst scan with immune-associated genes highlighted (green) and labeled were significant. Blue line: 95th-percentile empirical threshold; red line: 99th-percentile empirical threshold. IRF2, IL2RB, TNFSF18, TNFRSF19, and IL15 are labeled at their respective peak markers.

**Table 1.** Novel immune-associated genes identified in the buffalo-native (UOA_WB_1) iHS and Fst selection scans, ranked by strength of independent supporting evidence (KEGG pathway membership, EHH decay pattern, haplotype furcation structure, and the number of haplotypes carrying each allele at the focal marker). Tier 1 (Strong): broad, smooth EHH decay; well-resolved furcation structure on both allelic clades; balanced haplotype counts (minor allele n≥10/30). Tier 2 (Moderate): reasonable EHH and furcation evidence with somewhat asymmetric or shallower branching. Tier 3 (Suggestive): visually striking signal but supported by very few haplotypes carrying the minor allele (n≤5/30), raising the possibility that the extended-homozygosity pattern reflects a small-sample artifact (shared identity-by-descent among few individuals) rather than genuine positive selection; these findings should be interpreted with caution pending replication in a larger sample. None of the 14 classical bTB candidate genes examined (SLC11A1, BOLA-DRB3, TLR1, TLR2, TLR4, TLR6, IFNG, NOS2, SP110, VDR, GBP2, GBP6, CXCR1, CXCR2) reached the significance thresholds applied in either scan. KEGG pathway assignment used the cattle (Bos taurus) ortholog database (organism code ‘bta’) in the absence of a dedicated Bubalus bubalis pathway resource.

| Gene | Category | KEGG Pathway | Chr | Test (Evidence) | Haplotype n (Major/Minor) | EHH Signal | Furcation Structure | Confidence Tier | Putative Functional Role |
| --- | --- | --- | --- | --- | --- | --- | --- | --- | --- |
| *TNFSF18* | Novel immune-associated | Cytokine-cytokine receptor interaction (map04060) | 5 | Fst | 16 / 14 | Broad, smooth, symmetric decay | Strong, well-resolved both clades | Tier 1 - Strong | TNF-family ligand (GITRL); pairs with GITR, T-cell co-stimulation/regulation |
| *IL2RB* | Novel immune-associated | Cytokine-cytokine receptor interaction (map04060) | 4 | Fst | 20 / 10 | Moderate, asymmetric decay | Strong, clear branching both clades | Tier 1 - Strong | Shared receptor subunit for IL-2 and IL-15 signaling (γ-chain family) |
| *TNFRSF19* | Novel immune-associated | Cytokine-cytokine receptor interaction (map04060) | 13 | Fst | 16 / 14 | Broad, reasonably smooth | Decent, less depth on minor clade | Tier 2 - Moderate | TNF-receptor superfamily member (TROY); ligand unconfirmed, likely non-immune (neuronal/developmental) role |
| *IRF2* | Novel immune-associated | Type I interferon regulation (general) | 1 | Fst | 19 / 11 | Moderate decay | Real branching, both clades resolved | Tier 2 - Moderate | Transcriptional regulator of interferon-stimulated gene expression |
| *IL15* | Novel immune-associated | Cytokine-cytokine receptor interaction (map04060) | 17 | Fst | 25 / 5 | Weak, fast decay | Sparse minor-allele clade | Tier 3 - Suggestive | Cytokine signaling via IL15RA/IL2RB/IL2RG complex; T- and NK-cell homeostasis |
| *CD28* | Novel<br>immune-associated | T cell receptor<br>signaling<br>pathway<br>(map04660) | 2 | iHS | 27 / 3 | Dramatic<br>plateau<br>(minor allele) | Minor clade<br>nearly<br>absent (n=3) | Tier 3 -<br>Suggestive | Co-stimulatory receptor<br>driving T-cell activation and<br>downstream IL-2 production |

Orthology-based KEGG annotation using the *Bos taurus* reference indicated that four of the six genes participate in immune-related signalling pathways. *CD28* was assigned to the T-cell receptor signalling pathway (KEGG map04660), where it functions as a co-stimulatory receptor. Interaction between CD28 and its B7-family ligands enhances T-cell activation, proliferation, and cytokine production, including interleukin-2 secretion (Kanehisa et al., 2023; Sharpe, 2009). *IL2RB* and *IL15* were both assigned to the cytokine–cytokine receptor interaction pathway (KEGG map04060). IL2RB forms part of the shared IL-2/IL-15 receptor signalling system, together with the common gamma chain IL2RG, while IL-15 contributes to the maintenance and activation of natural-killer cells and memory CD8-positive T cells (Fehniger & Caligiuri, 2001; Waldmann, 2006).

*TNFSF18*, which encodes the glucocorticoid-induced TNFR-related ligand (GITRL), was also associated with the cytokine–cytokine receptor interaction pathway. Its known receptor is GITR, encoded by *TNFRSF18*, and engagement of the GITRL–GITR system can regulate effector T-cell activation, regulatory T-cell activity, inflammatory responses, and immune-cell survival (Nocentini et al., 2007; Ward-Kavanagh et al., 2016). Although *TNFRSF19* belongs to the tumour-necrosis-factor receptor superfamily, its annotation did not establish a confirmed ligand pair within the canonical KEGG pathway used in this study. Available evidence suggests that TNFRSF19 has important roles in neural development, epithelial biology, and stem-cell regulation, with its contribution to immune regulation remaining less clearly defined than that of other TNFRSF members (Hu et al., 1999; NCBI, 2026).

Finally, *IRF2* encodes a member of the interferon-regulatory-factor family. IRF2 acts as a transcriptional regulator of interferon-responsive genes and can modulate the balance between type-I-interferon activation and repression depending on cellular context (Honda et al., 2005; Tamura et al., 2008; Wang et al., 2024). Taken together, the six loci represent biologically plausible immune-related candidates, but their identification through a single genome-wide statistic and their annotation through a cattle orthology proxy mean that they should be interpreted as putative buffalo immune-associated genes, rather than confirmed bTB-resistance genes. Functional validation using buffalo-specific expression, genotype–phenotype association, and experimental immune-response data will be required to determine their role in resistance or susceptibility to *Mycobacterium bovis*.

### 3.5 Haplotype-based robustness assessment

Extended haplotype homozygosity (EHH) decay and haplotype-furcation patterns were examined for all six novel immune-associated genes to assess whether their genome-wide signals were supported by local haplotype structure (Figures 4–5). For each focal SNP, the number of phased haplotypes carrying the major and minor alleles was also recorded (Table 1). EHH measures the persistence of haplotype homozygosity as the distance from a focal marker increases, whereas furcation diagrams provide a more detailed view of how shared haplotypes break down on either side of that marker (Sabeti et al., 2002; Gautier & Vitalis, 2012; Gautier et al., 2017).

**Figure 4.**
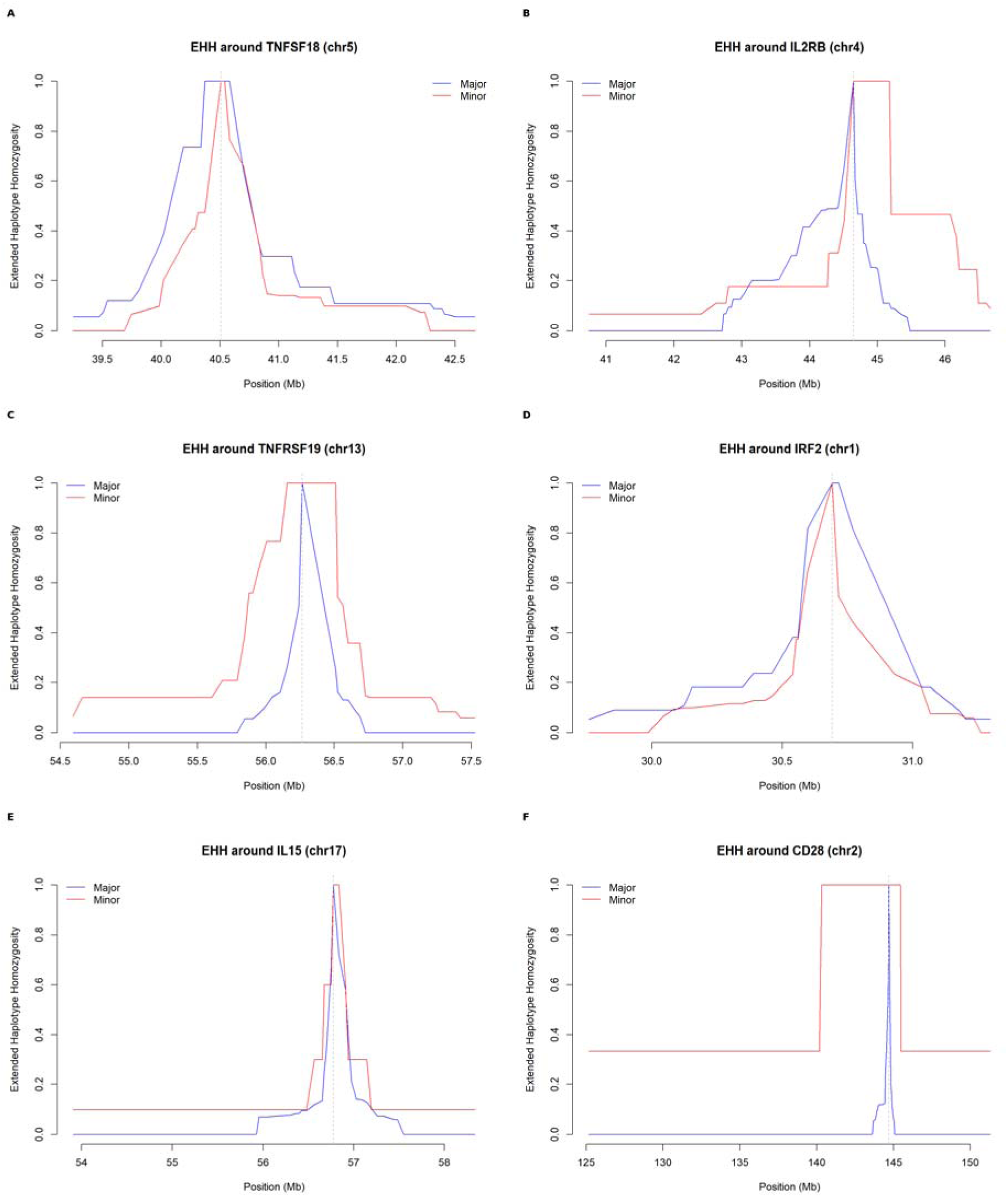
Extended haplotype homozygosity (EHH) decay around the peak SNP marker for each of the six novel immune-associated genes. (A) TNFSF18; (B) IL2RB; (C) TNFRSF19; (D) IRF2; (E) IL15; (F) CD28. Blue: major allele; red: minor allele. Curves are labeled “Major/Minor” rather than “Ancestral/Derived” because ancestral allele polarization is not available for Bubalus bubalis. Note the step-like, discrete decay in several panels (particularly E, F), a feature consistent with the limited haplotype sample size (n = 30) rather than a plotting artifact.

**Figure 5.**
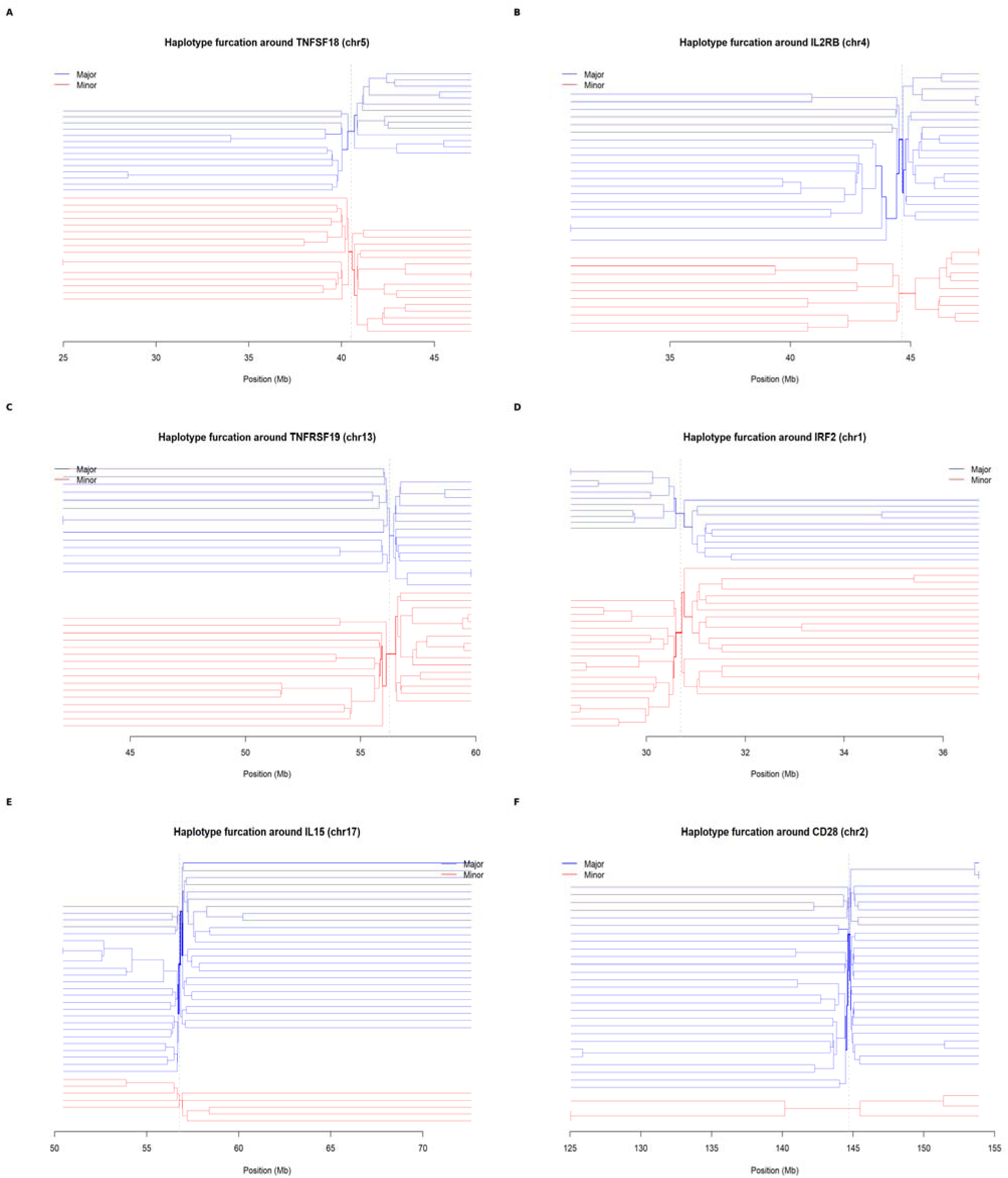
Haplotype furcation diagrams centered on the peak SNP marker for each of the six novel immune-associated genes. (A) TNFSF18; (B) IL2RB; (C) TNFRSF19; (D) IRF2; (E) IL15; (F) CD28. Blue clade: major allele; red clade: minor allele. Note the near-absence of internal branching in the minor-allele (red) clade for CD28 (F), consistent with this signal being carried by very few haplotypes (n = 3 of 30; Table 1).

*TNFSF18* and *IL2RB* showed the strongest concordant support. The focal markers had major-to-minor haplotype counts of 16:14 and 20:10, respectively. Both genes displayed extended and relatively smooth EHH decay, together with clearly resolved furcation structures for the two allele-specific haplotype groups. The combination of strong local haplotype sharing and substantial representation of the minor allele was consistent with a genuine extended haplotype signal rather than an artefact driven by only a few chromosomes.

*TNFRSF19* and *IRF2* provided intermediate support, with major-to-minor haplotype counts of 16:14 and 19:11, respectively. Their EHH profiles were compatible with extended haplotype structure, but the furcation trees showed less extensive or less sharply defined branching than those observed for *TNFSF18* and *IL2RB*. These loci were therefore classified as having moderate haplotype-based support.

The signals at *CD28* and *IL15* were interpreted more cautiously. Although both loci produced visually prominent genome-wide signals, their minor alleles were carried by only 3 and 5 of the 30 phased haplotypes, respectively. In particular, the *CD28* furcation diagram showed an almost absent minor-allele branch (Figure 5F). Such a pattern can arise when a long haplotype is shared by a small number of related or recently connected chromosomes, and does not necessarily indicate a population-wide selective sweep (Sabeti et al., 2002; Ralph & Coop, 2010). Accordingly, *CD28* and *IL15* were classified as suggestive rather than strong selection candidates.

Overall, the haplotype-based assessment provided the strongest independent support for *TNFSF18* and *IL2RB* (Appendix Figure 2 and Appendix 3), moderate support for *TNFRSF19* and *IRF2*, and more limited evidence for *CD28* and *IL15*. Because the haplotypes were analyzed without ancestral-allele polarization, the results are reported in terms of major and minor alleles rather than ancestral and derived alleles. These classifications were used as qualitative robustness categories and should not be interpreted as formal statistical significance levels.

## 4. Discussion

### 4.1 Main findings

This study used buffalo-native genomic coordinates to investigate selection signals and case–control differentiation in Nili-Ravi buffalo. The analysis identified 135 candidate iHS outliers and detected several markers with elevated *F*_ST_values. A subsequent immune-gene screen identified six genes outside the predefined classical bTB candidate panel: *TNFSF18, IL2RB, TNFRSF19, IRF2, IL15,* and *CD28*. These genes were detected through either the iHS or *F*_ST_analysis, but none was supported by both statistics.

The six loci should therefore be regarded as discovery-stage candidates rather than confirmed genes for resistance to bovine tuberculosis. Their biological interpretation was strengthened by three additional observations: several mapped to conserved immune-signalling pathways, the strongest candidates showed extended haplotype structure, and their functions were consistent with cellular immune responses known to contribute to control of *Mycobacterium bovis*. However, the small sample size, the limited marker density of the array, and the use of cattle orthologs for pathway annotation require cautious interpretation.

### 4.2 Classical bTB candidate genes

None of the 14 classical bTB candidate genes exceeded the predefined iHS threshold of—log_10_(p_iHS_) ≥ 4or the upper 5% *F*_ST_threshold. This result does not demonstrate that these genes are irrelevant to bTB resistance. The panel was assembled mainly from previous cattle and buffalo studies, whereas the present analysis tested unusually long haplotypes and allele-frequency differentiation. A gene may influence disease through modest-effect variants, expression differences, regulatory alleles, structural variants, or gene-by-environment interactions without producing an extreme genome-wide statistic.

The classical genes represent several important immunological mechanisms. *SLC11A1* encodes NRAMP1, a macrophage transporter that regulates metal availability within the phagosome and may restrict intracellular mycobacterial replication. *NOS2* encodes inducible nitric-oxide synthase, which contributes to macrophage-mediated killing of intracellular pathogens. The Toll-like receptors *TLR1, TLR2, TLR4,* and *TLR6* recognize microbial components and initiate inflammatory signalling through adaptor proteins such as MYD88. Buffalo studies have specifically examined *TLR* polymorphisms in relation to tuberculosis infection, providing species-relevant support for this pathway.

*BoLA-DRB3* is involved in antigen presentation to CD4-positive T cells, and its high polymorphism may alter the repertoire of mycobacterial peptides presented to the adaptive immune system. *IFNG* encodes interferon-γ, a central cytokine for activating infected macrophages and promoting Th1-type immunity. In cattle, IFN-γ release is widely used to measure cellular responses to *M. bovis*, and infected animals can generate antigen-specific T cells producing IFN-γ, IL-2, and TNF-α. *SP110* has been associated with intracellular pathogen control, whereas *VDR* may influence macrophage activation and antimicrobial gene expression. Finally, *CXCR1* and *CXCR2* regulate neutrophil recruitment, while *GBP2* and *GBP6* belong to the interferon-responsive guanylate-binding-protein family.

The lack of significant signals at these loci may reflect limited statistical power rather than the absence of biological effects. With only 15 animals, allele-frequency estimates are highly sensitive to individual chromosomes, and rare or low-frequency alleles may not be reliably evaluated. In addition, the 90K SNP array does not capture all buffalo-specific variants. Future studies should report nominal iHS and *F*_ST_values for the complete candidate panel, rather than reporting only whether each gene crossed the threshold. This would allow direct comparison across populations and facilitate meta-analysis as larger buffalo datasets become available.

### 4.3 Novel immune-associated genes

The unbiased screen identified a different group of genes that points toward adaptive immune regulation. *CD28, IL2RB, IL15,* and *TNFSF18* form the most coherent functional group, linking T-cell costimulation, cytokine signalling, lymphocyte expansion, and immune-cell survival.

*CD28* encodes a major T-cell costimulatory receptor. Its interaction with CD80 and CD86 on antigen-presenting cells strengthens T-cell receptor signalling, promotes IL-2 production, supports clonal expansion, and improves lymphocyte survival. These processes are relevant to bTB because effective control of intracellular mycobacteria depends on the development and maintenance of antigen-specific T-cell responses. In cattle, CD4-positive T-cell and IFN-γ responses are important components of immunity to *M. bovis*, while mouse models have demonstrated the importance of coordinated IL-12, IFN-γ, and T-cell responses for protection against tuberculosis (Cooper et al., 1993; Flynn et al., 1993; Waters et al., 2009).

*IL2RB*, also known as CD122, encodes the β-chain shared by the IL-2 and IL-15 receptor systems. It participates in JAK–STAT and PI3K–AKT signalling and contributes to the proliferation and survival of activated T cells, memory CD8-positive T cells, regulatory T cells, and natural-killer cells. Human and mouse studies show that impaired IL2RB signalling can disrupt lymphocyte development, regulatory T-cell function, and CD8-positive T-cell homeostasis. In the present study, the *IL2RB* signal is particularly interesting because it connects the costimulatory function of CD28 with cytokine-dependent lymphocyte expansion.

*IL15* supports natural-killer-cell development and the maintenance of memory CD8-positive T cells. It may help sustain cellular immune memory after exposure to mycobacterial antigens and may act through the shared IL2RB/IL2RG receptor components. Its identification does not establish a direct association with bTB resistance, but it provides a biologically plausible link between the detected genomic signal and the persistence of antimycobacterial cellular immunity.

*TNFSF18* encodes GITRL, a ligand of the TNF superfamily that binds TNFRSF18/GITR. GITR–GITRL signalling can regulate effector T-cell activation, regulatory T-cell function, and inflammatory responses. The identification of *TNFSF18* alongside *CD28, IL2RB,* and *IL15* suggests that selection may have affected a broader T-cell regulatory network rather than a single isolated immune gene.

The remaining genes require more cautious interpretation. *IRF2* encodes an interferon-regulatory transcription factor that can modulate type-I-interferon-responsive genes. It may influence the balance between effective antimicrobial signalling and excessive inflammation, but its role in buffalo bTB resistance remains uncertain. *TNFRSF19*, also known as TROY, is structurally a member of the TNF-receptor superfamily, but its best-characterized functions in mammals involve neural development, epithelial biology, and stem-cell regulation. Its detection may reflect a genuine but poorly characterized immune function, a regulatory effect on a nearby gene, or nearest-gene assignment rather than direct causality.

The cattle and buffalo literature supports a polygenic interpretation of bTB resistance. Genome-wide studies in cattle have identified multiple resistance-associated regions rather than a single universal locus, and comparative analyses of zebu and taurine cattle have implied diverse immune and non-immune genes. Differential immune-gene expressions have also been observed between buffalo and cattle following infection, supporting the possibility that species and breed differences in immune regulation contribute to disease outcome.

### 4.4 Haplotype-based support

The haplotype analysis demonstrated that the six novel genes did not have equal evidentiary support. *TNFSF18* and *IL2RB* showed the strongest concordance among EHH decay, furcation structure, and allele-specific haplotype counts. Their minor alleles were carried by 14 and 10 of the 30 phased haplotypes, respectively. This relatively balanced representation makes it less likely that their extended haplotype patterns were generated solely by one or two related chromosomes.

*TNFRSF19* and *IRF2* showed moderate support. Their allele counts were reasonably balanced, but their furcation structures were less distinct and their EHH profiles were less consistent than those of *TNFSF18* and *IL2RB*. These loci should be retained for replication but not assigned the same confidence as the two stronger candidates.

The *CD28* and *IL15* signals were more fragile. Their minor alleles occurred on only 3 and 5 of the 30 phased haplotypes, respectively. The *CD28* furcation diagram contained almost no minor-allele branch, suggesting that the striking signal may have been driven by a small number of shared chromosomes. Extended haplotype homozygosity reflects local haplotype sharing, and such sharing can arise from recent common ancestry or identity-by-descent as well as selection. Consequently, a long haplotype in a few individuals should not automatically be interpreted as a population-wide selective sweep.

The tiered classification used here—strong, moderate, and suggestive—was therefore an important safeguard. It prevented all markers exceeding a numerical threshold from being treated as equally reliable. Nevertheless, the tiers are qualitative and should be validated using larger cohorts, simulations, permutation tests, and independent populations.

### 4.5 Importance of buffalo-native coordinates

The coordinate-remapping analysis produced very similar global *F*_ST_values for the original cattle-proxy and buffalo-native datasets. The mean and weighted estimates were 0.0099 and 0.0157 using the bovine-proxy coordinates, compared with 0.00945 and 0.0156 after remapping. This agreement indicates that the broad case–control differentiation pattern was stable and that the coordinate conversion did not materially alter the genome-wide population-genetic results.

However, similar global statistics do not guarantee identical biological interpretation. The gene assigned to a significant SNP depends on its chromosome, physical position, gene model, and surrounding annotation. A marker may appear close to one gene under a cattle-based system but map to a different genomic context in buffalo. Correct species-specific coordinates are therefore essential for gene annotation, pathway mapping, and interpretation of regulatory regions.

The chromosome-level water buffalo assembly provides an appropriate framework for this analysis because river buffalo has 25 chromosome pairs, whereas cattle-based resources use a different chromosome organization. The updated buffalo array map developed by Nascimento et al. (2021) enables existing array datasets to be remapped without discarding valuable historical genotypes. As buffalo-specific annotations and breed-matched assemblies improve, buffalo-native reanalysis should become standard for selection scans and association studies.

### 4.6 Limitations and future directions

The principal limitation of this study was the small sample size. With 85 animals, the number of independent haplotypes is limited, allele-frequency estimates are imprecise, and long haplotypes may be disproportionately influenced by relatedness or recent common ancestry. The F_ST_comparison may also reflect sampling structure, family relationships, or differences in disease classification in addition to genetic differentiation.

The 90K array represents another limitation. Array markers may not tag rare buffalo-specific alleles, structural variants, copy-number changes, or regulatory variants that contribute to bTB resistance. Whole-genome sequencing would provide improved resolution around the six candidate loci and would allow examination of local linkage disequilibrium, rare variants, structural variation, and regulatory elements.

The use of cattle orthologs for KEGG and DAVID annotation was necessary because buffalo annotation coverage was limited, but it introduces uncertainty. Orthology-based pathway assignments should therefore be treated as conserved functional predictions rather than buffalo-specific evidence. Future analyses should incorporate buffalo gene models, expression atlases, regulatory annotations, and experimentally validated immune pathways.

The six genes should next be tested in larger Nili-Ravi cohorts and independent buffalo populations. RNA sequencing or targeted expression assays in macrophages, lymphocytes, and infected tissues could determine whether *CD28, IL2RB, IL15,* and *TNFSF18* respond to *M. bovis* exposure. Functional assays measuring T-cell proliferation, IFN-γ and IL-2 production, IL-15 responsiveness, STAT5 activation, macrophage antimicrobial activity, and GITR–GITRL signalling would provide a direct test of the proposed mechanisms. Such work would establish whether these signals represent genuine buffalo adaptations to mycobacterial challenge or population-specific haplotype patterns requiring alternative explanations.

## 6. Conclusion

Reanalyzing Axiom Buffalo 90K genotype data from 85 Nili-Ravi buffalo against buffalo-native genomic coordinates (UOA_WB_1) revealed no evidence of selection across 14 classical, cattle-derived bTB candidate genes. Instead, an unbiased genome-wide screen uncovered six candidate genes enriched for immune function. Four of these locus signatures converge on T-cell stimulation and cytokine receptor signaling, with two genes TNFSF18 and IL2RB validated across three independent, haplotype-based selection metrics. These findings indicate that adaptive, cell-mediated immune pathways may play an underappreciated role in host genetic resistance to bTB in Nili-Ravi buffalo. Furthermore, this study highlights the necessity of using species-specific reference coordinates combined with multi-method haplotype validation to ensure accurate, robust selection-scan inferences in historically under-resourced livestock species.

## Author Contributions

H.M. conceived and designed the study. A.A. and A.B. coordinated data acquisition and assisted with dataset preparation. H.M. performed data curation, quality control, LD decay, and Ne analyses and led the bioinformatics workflow. W.A.K. provided technical support for genomic analyses and contributed to data interpretation. S.M.L. and A.M. facilitated sample logistics and contributed domain expertise related to buffalo breeding systems. S.M.L. contributed to population-level context and breed interpretation, while A.M. supported institutional coordination and data validation. A.A. contributed to sample facilitation and provided logistical support through Maxim Agri (Pvt.) Ltd. H.M. and H.K. carried out candidate gene annotation, interpreted the results, and drafted the manuscript. All authors critically reviewed the manuscript, provided intellectual input, and approved the final version for submission.

## Institutional Review Board Statement

NA

## Data Availability Statement

The generated data of this study is available with the author and available upon a reasonable request.

## Acknowledgments

The authors acknowledge the Higher Education Commission (HEC) of Pakistan for facilitating access to genotype data. We thank the Livestock and Dairy Development Department (L&DD), Punjab, and other collaborating institutions for their support in sample collection and data generation. The authors also appreciate the technical support provided by colleagues at the Department of Animal Breeding and Genetics, University of Veterinary and Animal Sciences (UVAS), Lahore. Maxim Agri (Pvt.) Ltd. is gratefully acknowledged for providing samples and logistical support.

## Appendix -I

| Gene | Relevance and supporting evidence |
| --- | --- |
| *SLC11A1* | Encodes NRAMP1, a macrophage metal-transporter involved in intracellular control of mycobacteria. Polymorphisms in cattle have been investigated in relation to resistance to bovine tuberculosis and other mycobacterial diseases (Allen et al., 2010; Kadarmideen et al., 2011). |
| *BoLA-DRB3* | A highly polymorphic bovine major-histocompatibility-complex class II gene involved in antigen presentation. *BoLA-DRB3* variation has been linked to differences in immune responses to *M. bovis* antigens in cattle (Allen et al., 2010). The gene is also polymorphic in buffalo, supporting its use as a comparative immune-response candidate (Miretti et al., 2001; Eltarabany et al., 2018). |
| *TLR1* | Part of the Toll-like-receptor system that recognizes microbial lipoproteins and initiates innate immune signalling. Polymorphisms in *TLR1* have been evaluated for associations with tuberculosis susceptibility in Chinese Holstein cattle (Sun et al., 2012). |
| *TLR2* | Recognizes mycobacterial cell-wall components and contributes to macrophage activation. *TLR2* has been identified as a biologically relevant tuberculosis-resistance candidate in cattle and other mammalian hosts (Allen et al., 2010). |
| *TLR4* | Participates in innate immune recognition and inflammatory signalling. Bovine *TLR4* polymorphisms have been examined in studies of mycobacterial infection and related production-animal diseases, supporting its inclusion as an immune-response candidate (Allen et al., 2010; Muriuki et al., 2021). |
| *TLR6* | Forms heterodimers with *TLR2* and contributes to recognition of microbial lipoproteins. Its inclusion is supported by the broader tuberculosis literature and by the functional importance of the *TLR2/TLR1/TLR6* signalling axis in antimycobacterial immunity (Allen et al., 2010; Schurz et al., 2015). |
| *IFNG* | Encodes interferon-γ, a central cytokine for macrophage activation and cellular immunity against *M. bovis*. Variation in bovine *IFNG* has been investigated in relation to mycobacterial infection and interferon-γ production phenotypes in cattle (Allen et al., 2010; Muriuki et al., 2021). |
| *NOS2* | Encodes inducible nitric-oxide synthase, which contributes to macrophage-mediated antimicrobial activity. Polymorphisms in bovine *NOS2/NOS2A* have been associated with bTB susceptibility or resistance in Holstein cattle (Cheng et al., 2016; Chai et al., 2021). |
| *SP110* | Encodes a nuclear-body protein involved in intracellular pathogen control and macrophage responses. Functional studies demonstrated that *SP110* expression can reduce *M. bovis* replication and pathology in cattle (Wu et al., 2015). |
| *VDR* | Encodes the vitamin-D receptor, a transcriptional regulator involved in macrophage activation and antimicrobial immune responses. Its inclusion is biologically justified by the role of vitamin-D signalling in antimycobacterial immunity, although direct bTB-association evidence in cattle or buffalo is more limited than for *SLC11A1*, *NOS2*, or *SP110* (Allen et al., 2010). |
| *GBP2* | Encodes a guanylate-binding protein involved in interferon-responsive |
|  | antimicrobial and intracellular-pathogen pathways. The gene was retained as a mechanistically plausible candidate because guanylate-binding proteins are part of the host response to intracellular pathogens; direct buffalo-specific bTB association evidence remains limited (O’Grady et al., 2025). |
| *GBP6* | A member of the interferon-inducible guanylate-binding-protein family. Its potential relevance is based on the role of this gene family in intracellular pathogen defence and its occurrence among immune-response candidates in cattle genomic and transcriptomic studies; direct functional evidence for bTB in buffalo remains to be established (O’Grady et al., 2025). |
| *CXCR1* | Encodes a chemokine receptor involved in neutrophil migration and inflammatory-cell recruitment. Bovine *CXCR1* variation has been studied in relation to immune and health traits, and the receptor is biologically relevant to leukocyte trafficking during infection (Rinaldi et al., 2011). |
| *CXCR2* | Encodes a second major chemokine receptor involved in neutrophil recruitment and inflammatory signalling. Its inclusion is supported by the documented importance of the bovine chemokine-receptor system in host defence, although direct bTB-specific association evidence is currently limited (Rinaldi et al., 2011; Allen et al., 2010). |

**Appendix Figure 1:**
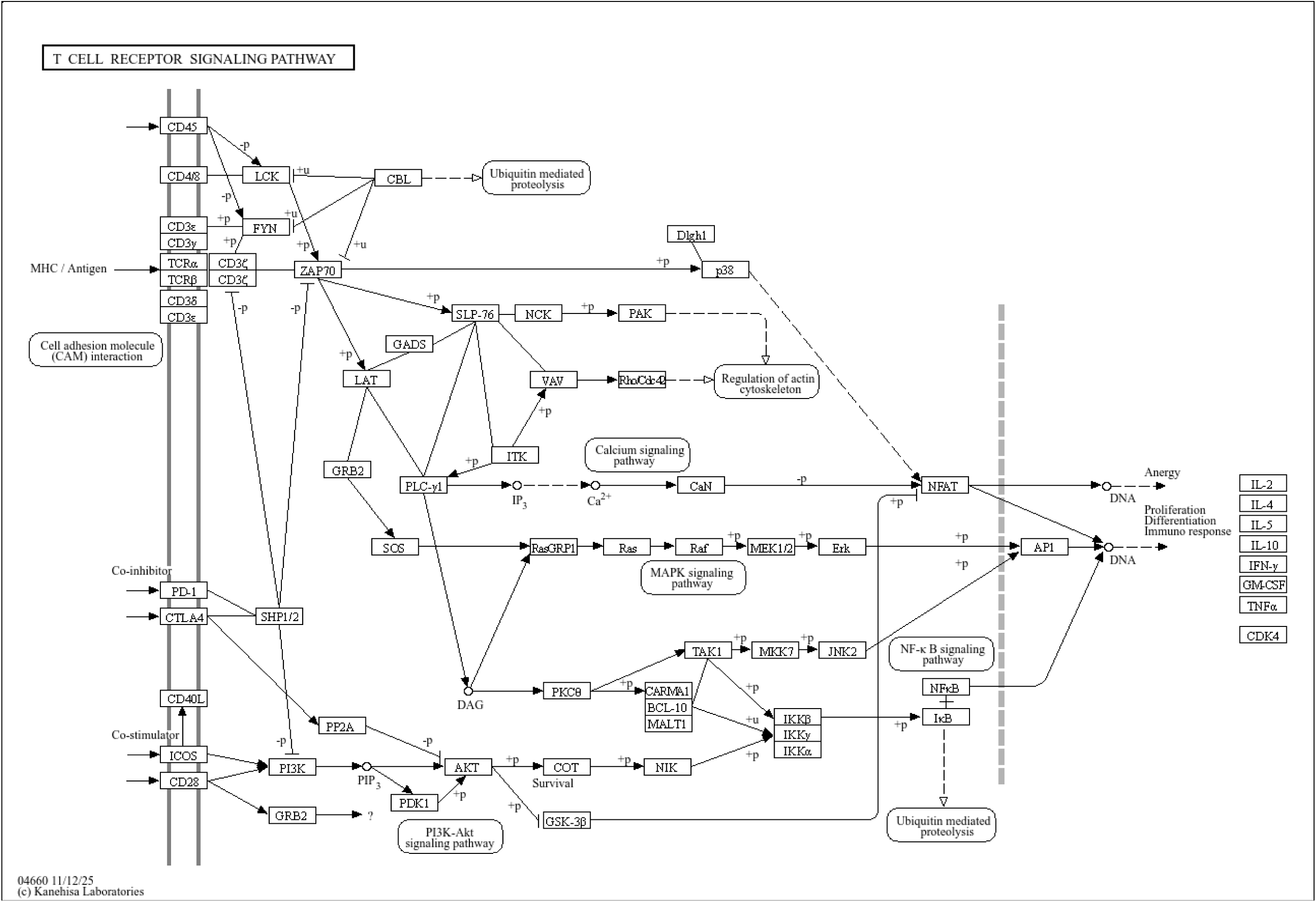
CD28 was assigned to the T-cell receptor signalling pathway (KEGG map04660), where it functions as a co-stimulatory receptor. Interaction between CD28 and its B7-family ligands enhances T-cell activation, proliferation, and cytokine production, including interleukin-2 secretion (Kanehisa et al., 2023; Sharpe, 2009).

**Appendix Figure 2:**
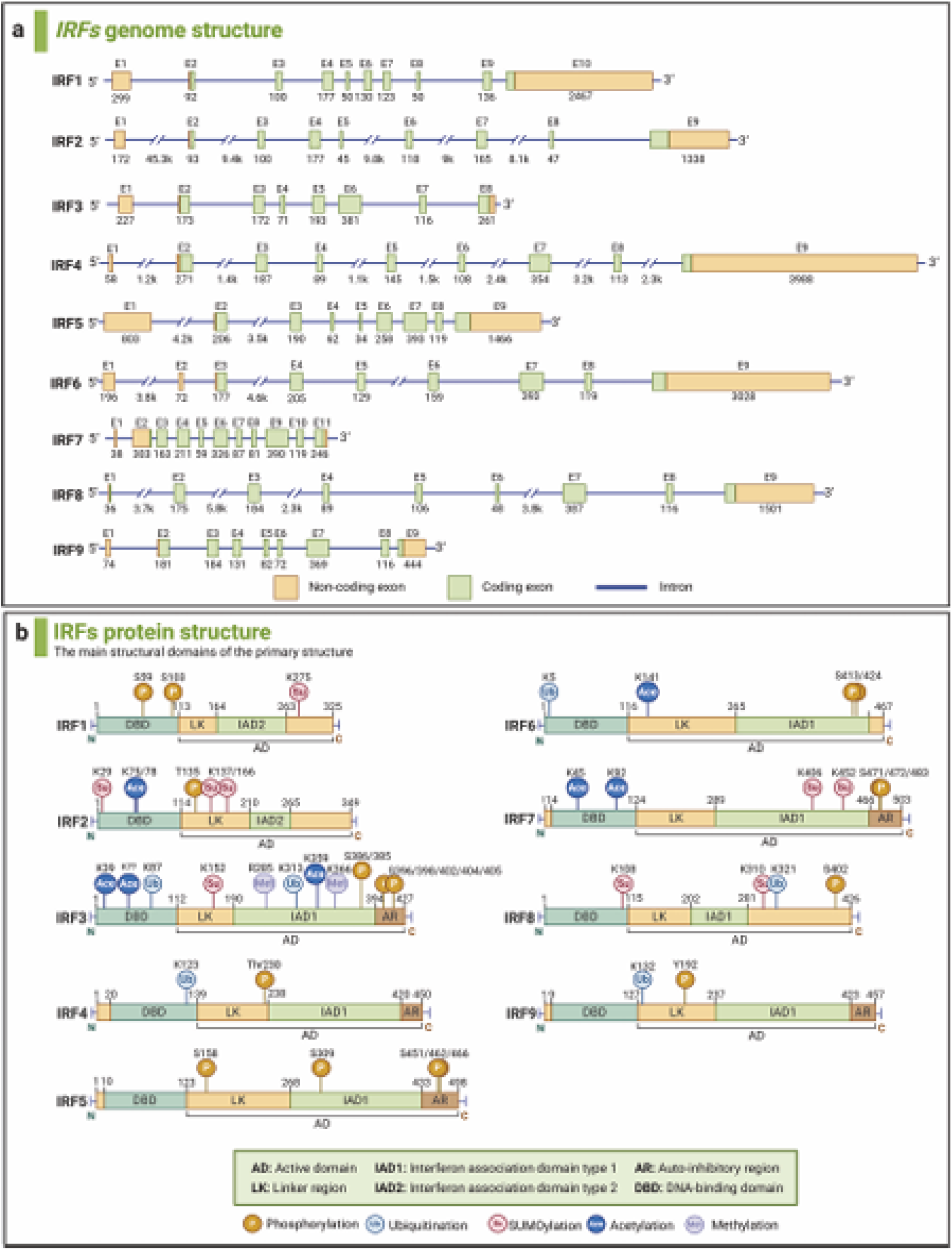
The genes and proteins structures of IRF family members. A) The genomic structures. The boxes mark the exons, including non-coding exons (orange) and coding exons (green). blue lines mark the introns. Start and stop codons are indicated. The numbers below the genes indicate the sizes of the exons and introns. B) The structures of IRF family proteins. DBD represents the DNA-binding domain, LK represents the linker region, IAD represents interferon association domain, and AR represents auto-inhibitory region. Active domains (AD) are also marked on the figure. The upper numbers refer to the starting amino acid sites of diferent domains, and the different post-translational modifications of the IRFs are presented, including Phosphorylation, Ubiquitylation, SUMOylation, Methylation, and Acetylation splicing variants (Wang et al., 2024).

**Appendix Figure 3:**
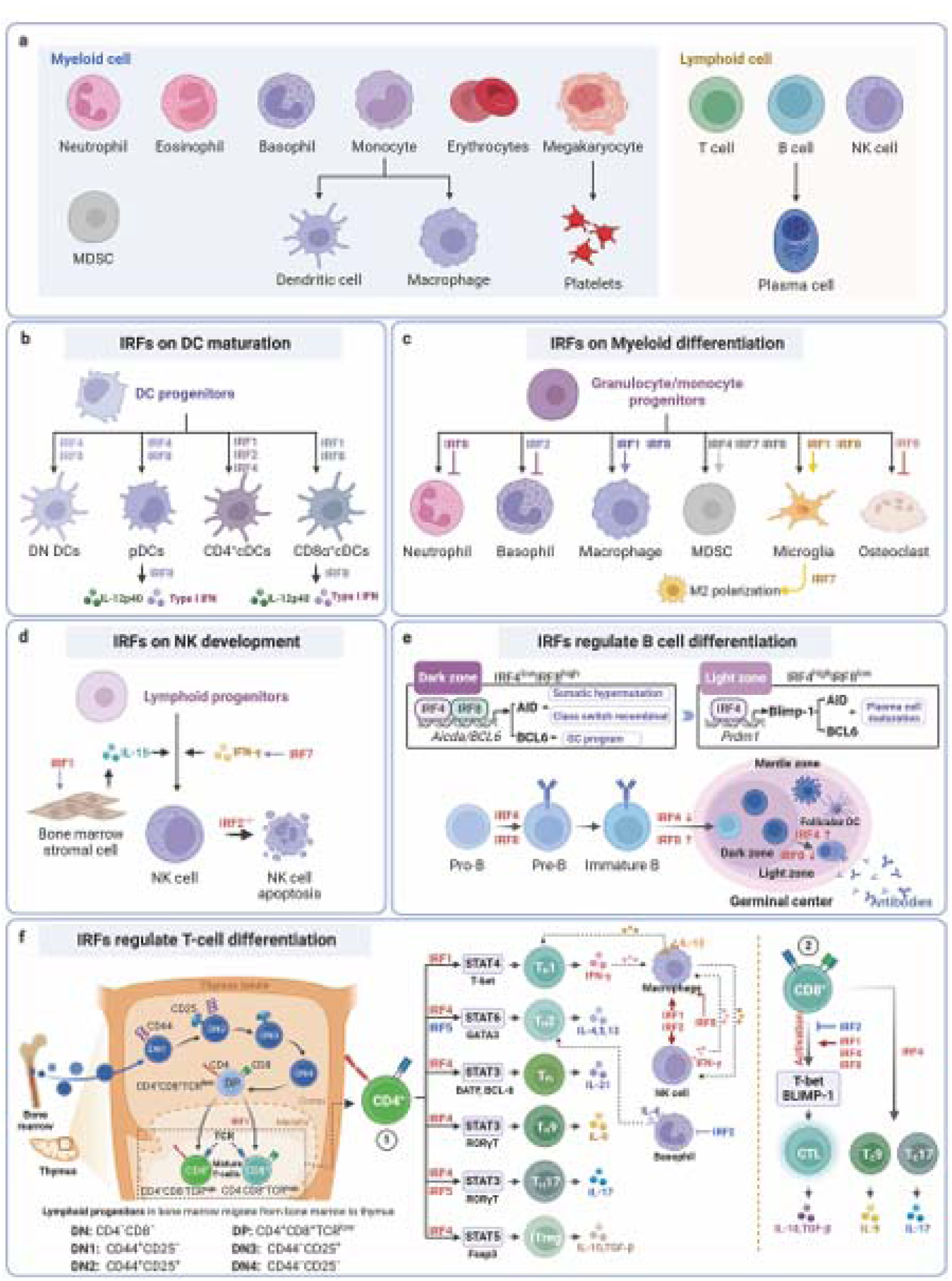
The regulatory effects and molecular mechanisms of IRFs on immune cell development. a The subtypes of myeloid cells and lymphoid cells. b The induction effect of IRFs on DC maturation and cytokine production. c The role of IRFs in myeloid differentiation and MDSC aggregation. d Regulation of IRFs in Natural Killer Cells development. e Multistep regulation of B cells and plasma cells by IRFs. f IRFs regulate T cell development and differentiation (Wang et al., 2024).

**Appendix Figure 4:**
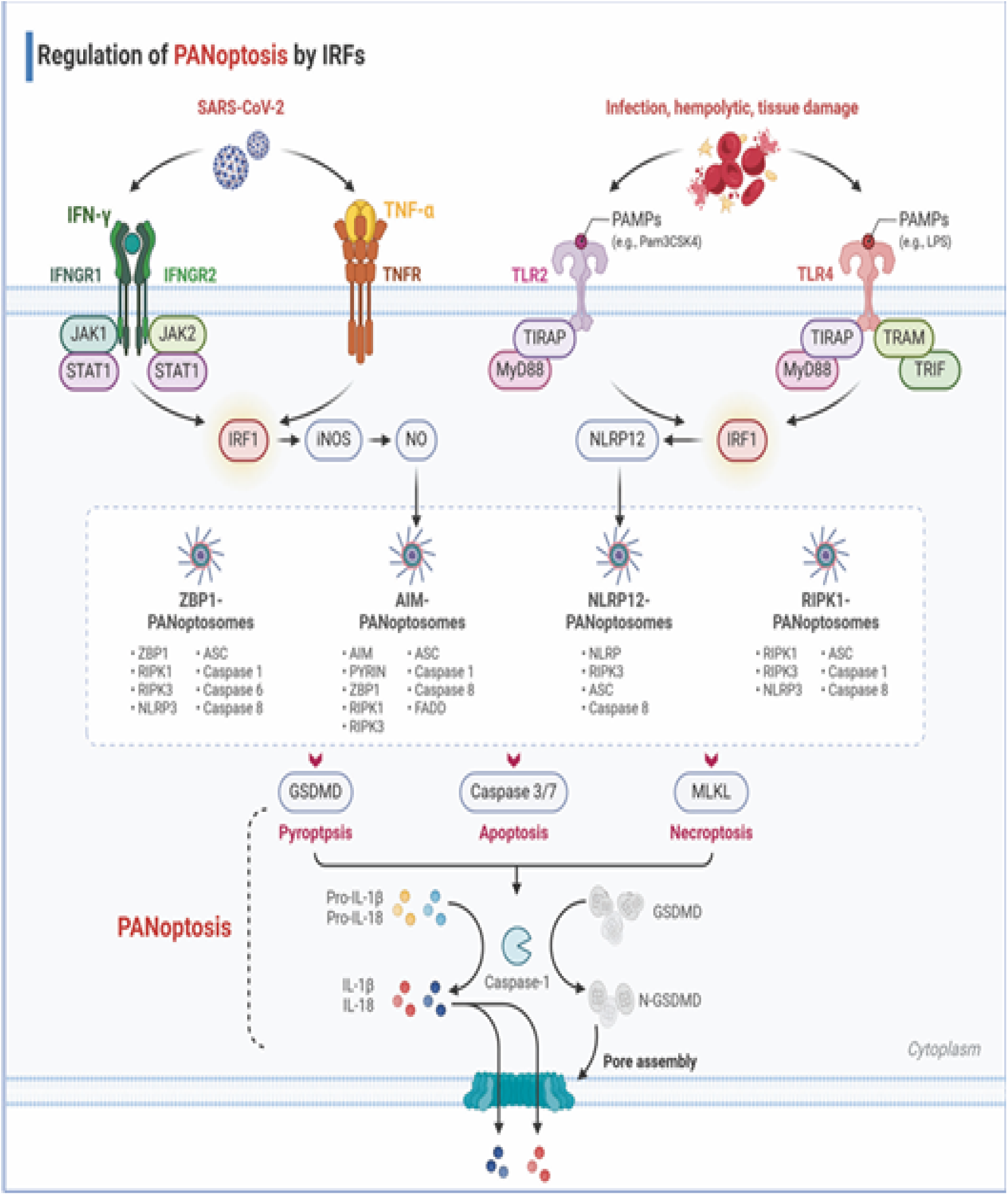
Regulation of apoptosis by IRFs. IRF-1 is a tumor suppressor and a regulatory factor of the IFN-γ system, with IFN-γ promoting the expression of IRF-1. IFN-α also induces the rapid phosphorylation and DNA-binding activity of STAT-1, followed by the accumulation of IRF-1 and IRF7. These two transcription factors bind to the TRAIL promoter and induce TRAIL expression. TRAIL is a key participant in the apoptotic pathway and plays a significant role in IFN-induced cell killing. IRF-3 is also involved in the transcriptional induction of TRAIL, where it transactivates the TRAIL promoter upon viral infection, upregulating TRAIL transcription. Conversely, IRF4 actively inhibits the transactivation mediated by IRF1. IRF3 triggers apoptosis through the RIPA pathway, which depends on the linear ubiquitination of specific lysine residues of IRF3. Within the RIPA signaling pathway, IRF3 interacts with the pro-apoptotic protein Bax to form the IRF3-Bax complex and translocates to the mitochondria, catalyzing the release of cytochrome C into the cytoplasm, subsequently activating Caspase, and ultimately leading to apoptosis. Otulin, a deubiquitinase that removes linear ubiquitin chains, inhibits RIPA by deubiquitinating IRF3 in virus-infected cells malignancies, IRF5 promotes cancer cell proliferation (Wang et al., 2024).

**Appendix Figure 5:**
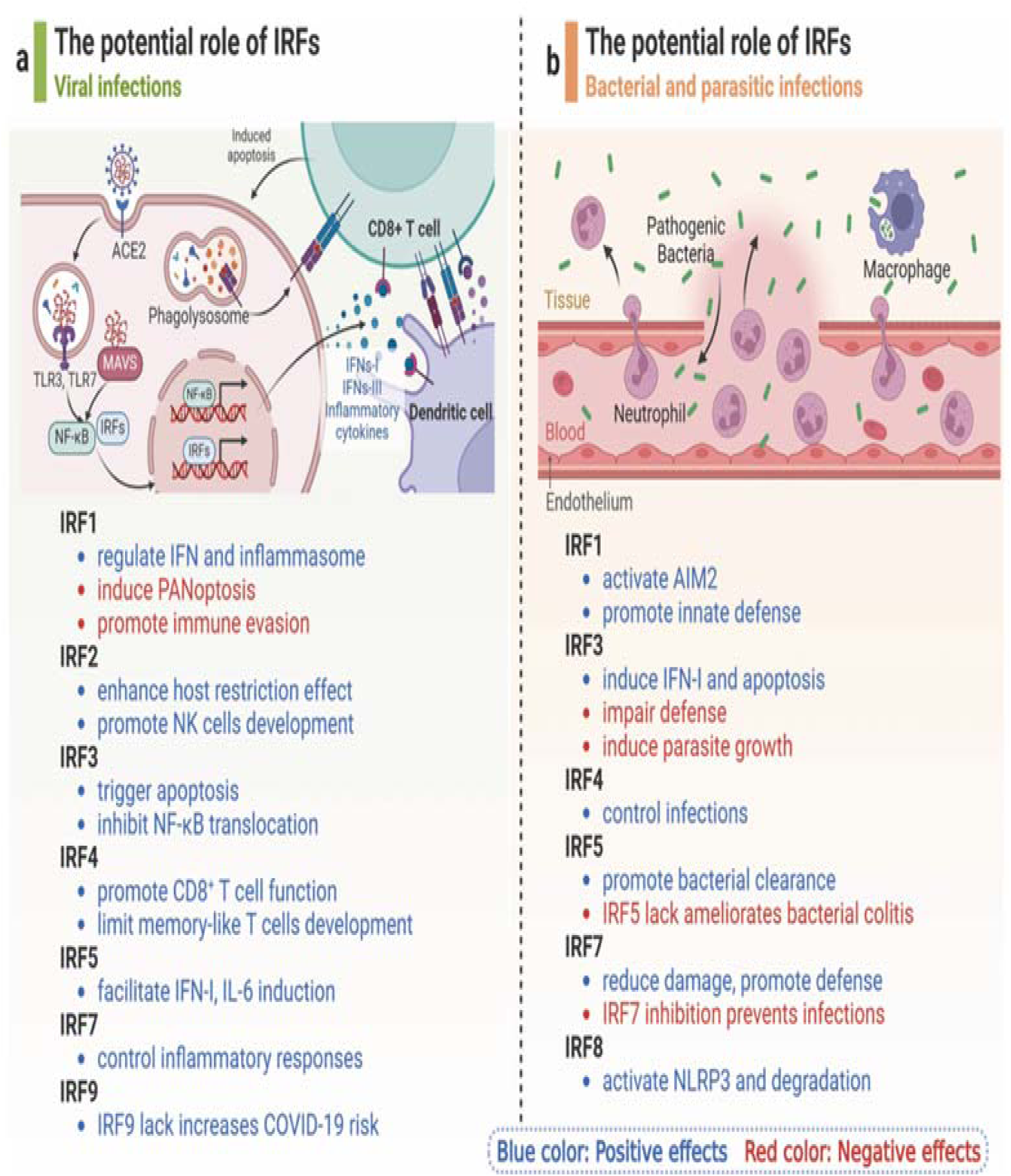
The potential role of IRFs in infectious diseases. a The roles of IRFs in viral infections. IRF1 can regulate IFN and inflammasome to fight viral infections, while it also induces cell PANoptosis and virus immune evasion. IRF2, IRF3, IRF4,IRF5, IRF7, and IRF9 play protective roles to prevent viral infections. b The roles of IRFs in bacterial and parasitic infections. IRF1, IRF4, and IRF8 prevent host aginst bacterial infections. IRF3, IRF5, and IRF7 are double-edged swords in bacterial and parasitic infections degradation in various cell types, which drive antiviral immunity to multiple clinically important viruses, including hepatitis C virus (HCV), West Nile virus,yellow fever virus, human immunodeficiency virus type 1(HIV1), gamma herpes virus, and human metapneumovirus, and VSV infection (Wang et al., 2024).

**Appendix Figure 6:**
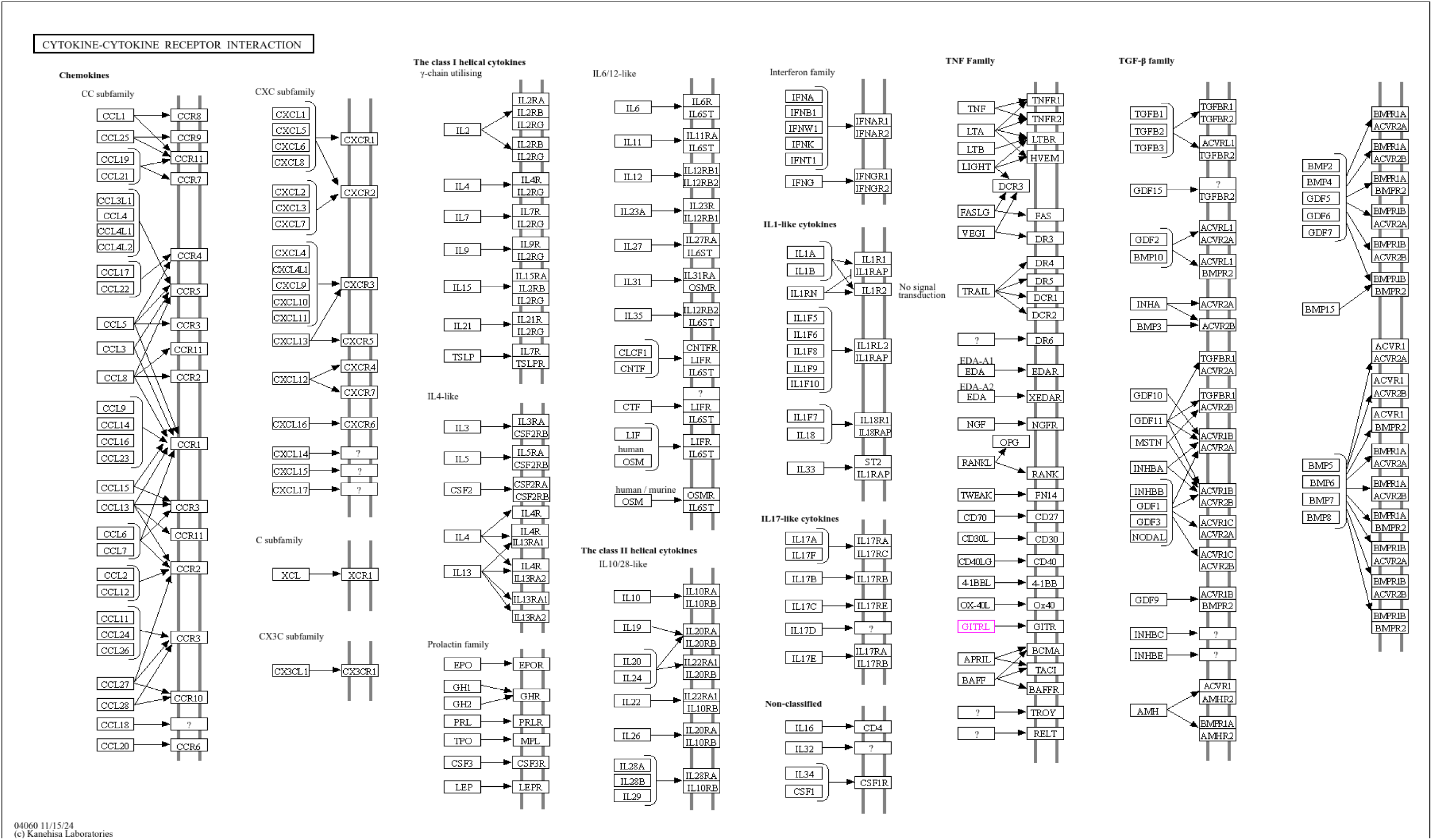
KEGG pathway annotation and immune function of **TNFSF18.** The buffalo candidate locus was mapped to the cattle ortholog TNFSF18 (GITRL), which corresponds to KEGG Orthology entry K05479. The gene was assigned to the cytokine–cytokine receptor interaction pathway (KEGG map04060) and the cytokine/neuropeptide BRITE hierarchy. TNFSF18 encodes a member of the tumour-necrosis-factor ligand superfamily and acts as the ligand for TNFRSF18, also known as GITR. The TNFSF18–TNFRSF18 interaction can regulate T-cell activation, proliferation, survival, and immune-response intensity, making this gene a biologically plausible immune-associated candidate (Gurney et al., 1999; Gaur & Aggarwal, 2003; Ward-Kavanagh et al., 2016). The pathway assignment was based on the cattle ortholog proxy in KEGG and should therefore be interpreted as a conserved functional annotation rather than direct experimental confirmation in water buffalo.

**Appendix Figure 7:**
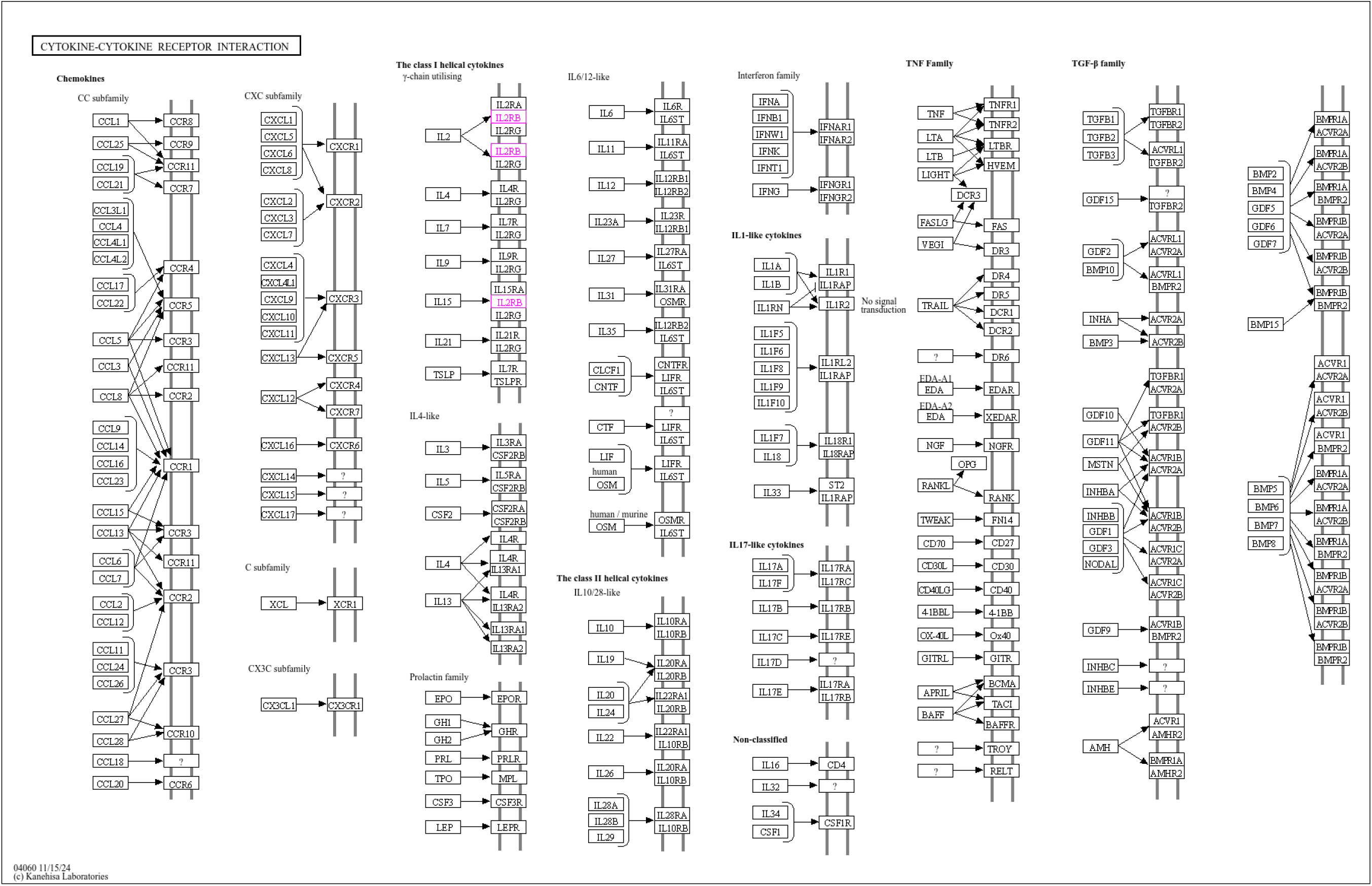
KEGG-based functional annotation of the **IL2RB** candidate gene. The candidate locus was assigned to KEGG Orthology group K05069 (IL2RB/CD122) and to the cytokine–cytokine receptor interaction, JAK–STAT, PI3K–Akt, Th1/Th2 differentiation, and Th17 differentiation pathways. IL2RB encodes the β-chain shared by the IL-2 and IL-15 receptor systems and contributes to cytokine-mediated signaling in T cells and natural killer cells. The annotation was obtained using the cattle ortholog proxy and is therefore presented as a comparative functional prediction for the buffalo genome.

